# Engineering a dynamic, controllable infectivity switch in bacteriophage T7

**DOI:** 10.1101/2021.08.23.457391

**Authors:** Chutikarn Chitboonthavisuk, Chun Huai Luo, Phil Huss, Mikayla Fernholz, Srivatsan Raman

**Author notes:** **Corresponding Author Srivatsan Raman –** *Department of Biochemistry, University of Wisconsin – Madison, 433 Babcock Dr, Madison, Wisconsin 53706, United States;. These authors contributed equally to this work (Chutikarn Chitboonthavisuk, Chun Huai Luo).

## Abstract

Transcriptional repressors play an important role in regulating phage genomes. Here, we examined how synthetic regulation based on repressors can be to create a dynamic, controllable infectivity switch in bacteriophage T7. We engineered T7 by replacing a large region of the early phage genome with combinations of ligand-responsive promoters and ribosome binding sites (RBS) designed to control the phage RNA polymerase. Phages with the engineered switch showed virulence comparable to wildtype when not repressed, indicating the phage can be engineered without a loss of fitness. When repressed, the most effective switch used a TetR promoter and a weak RBS, resulting in a two-fold increase in latent period (time to lyse host) and change in phage titer. Further, phage activity could be tuned by varying inducer concentrations. Our study provides a proof of concept for a simple circuit for user control over phage infectivity.

## Introduction

Bacteriophages (or ‘phages’) are obligate parasites which require a bacterial host to complete their life cycle^1^. Once a phage infects its host, a choreographed cascade of phage genes is expressed to regulate subsequent steps in the phage’s life cycle. Gene regulation in phages is frequently controlled by transcription repressors^2^. Transcription repressors act as switches that determine lifestyle decisions (e.g.: lytic vs. lysogenic) by silencing or activating different sets of genes. This is best exemplified in the transcription repressor-based genetic switch in phage lambda and other temperate phages with similar mechanisms (P22^3^, 434^4^, ϕC31^5^). Phage lambda persists in lysogenic state when transcription repressor CI represses early-stage lambda promoters, halting the transcriptional cascade of lytic genes. To activate lytic genes and release the prophage, a different repressor Cro, counteracts CI through differential binding at the same promoters. Transcription repressors also control activation and inactivation of non-integrating, episomal phage genomes, called pseudolysogens, in a nutrient dependent manner^6^. Transcription repressors are valuable tools for phages because their mechanism of gene regulation simply relies on steric obstruction of the RNA polymerase making it largely host-independent^7^. Further, a transcription repressor with strong affinity for its promoter can exert tight regulation over multiple open reading frames commonly found in long phage operons. While natural transcription regulation in phages is well studied, the engineering rules of introducing synthetic regulation into phages has not been developed.

Bacteria have long served as a popular chassis for exploring and prototyping synthetic regulation. This choice was largely driven by the availability of tools for bacterial genome engineering and the application goal of bacterial biomanufacturing^8^. In recent years, a strong impetus has emerged for engineering phages, driven similarly by the availability of new tools for phage genome engineering and their potential applications in medicine and biotechnology^9–11^. Phage genomes can be now edited with high precision from single base to kilobase resolution using yeast cloning, homology-directed repair, and enzymatic recombination^12,13^. Phages engineered using these approaches could be powerful tools for killing antibiotic resistant bacteria and precisely manipulating microbiomes with applications in agriculture, livestock, medicine, and environment. Engineered phages have many advantages over their natural counterparts including superior efficacy, greater programmability, higher compositional stability, and easier scalability of production^14^. Despite this promise, even basic rules of engineering new regulation into phages have not been developed. This is in stark contrast to the wealth of research on engineering promoters, switches, circuits, and pathways in bacteria. Rules for engineering bacteria are not directly transferrable to phages due to substantial differences in genome compactness, compartmentalization of regulation and kinetics of growth and replication. Thus, rules for engineering new regulation for phages must be developed by systematic design-build-test-learn analysis.

With this goal in mind, we examined how bacterial transcription repressors could be used to engineer synthetic genetic regulation in phages. We sought to engineer a repressor-based infectivity switch that can dynamically control the activity of an obligate lytic phage in a ligand-dependent manner. Such a system would provide direct user control of phage activity which would otherwise undergo unregulated exponential amplification upon infecting a host^15^. Natural phages such as lambda are not ligand-inducible but instead rely on stochastic differences in the concentration of CI vs. Cro to determine lysogenic vs. lytic choice^16^. A dynamic controllable infectivity switch would be a valuable tool for activating a phage at user-defined times to carry out microbiome editing in a natural or synthetic communities. Furthermore, inducible phage would be an effective form of biocontainment, as the phages only remain active while the ligand is provided but otherwise remain inert.

Here, we engineered an obligate lytic T7 phage with a synthetic regulatory switch to control its infection cycle in a ligand-dependent manner. To introduce synthetic regulation, we removed a large tract of native regulatory sequence from the phage genome and replaced this region with a short ligand-responsive bacterial promoter that regulates the T7 RNA polymerase gene. We saw no appreciable loss of fitness in the synthetic phage relative to wildtype T7 (T7_WT). We tested different ligand-responsive bacterial promoters and ribosome-binding sites (RBS) representing a range of expression levels to characterize how these variables affect phage activity. We measured our ability to attenuate phage replication by measuring the phage latent period and change in phage titer and found that the strongest attenuation of phage replication, or the ‘OFF’ state, occurred with a Tet-regulated promoter and a very weak RBS, which resulted in a 2-fold increase in latent period and an approximately 2-fold decrease in a change in phage titer. Ligand induced activation to the ‘ON’ state restored activity of synthetic phages to levels comparable to unregulated T7_WT phage. Our study provides a basic proof-of-concept for recoding a phage genome with synthetic regulation, paving the way of engineering more sophisticated circuitry to enable phages to carry out complex, user-defined tasks.

## Results

### Refactored synthetic phages retain wildtype infectivity

We sought to create a simple, dynamically controllable infectivity switch for phages using ligand-responsive transcriptional repressors naturally utilized for gene regulation in bacteria. To test our synthetic switch, we chose bacteriophage T7, a well characterized, prototypical obligate lytic phage that infects *Escherichia coli*. Bacterial systems are routinely engineered with inducible gene expression by placing ligand-responsive promoters upstream of gene of interest^17^. Bacterial genomes are generally tolerant to promoter substitutions as most substitutions have marginal effects on bacterial fitness^18^. In contrast, phage genomes have evolved to maintain compact genomes containing mostly essential genes with overlapping regulation^19–21^. This makes it challenging to identify a suitable T7 gene whose promoter can be replaced by ligand-responsive bacterial promoter.

The T7 phage genome is approximately 40 kb long and is roughly partitioned into early, middle and late-stage genes denoting their roles and when they are expressed at different stages of T7 lifecycle^22^. Transcription of phage genes occurs in two stages after infecting its host. First, the host *E. coli* RNA polymerase transcribes T7 RNA polymerase and other early phage genes^23–26^. After early genes have been expressed, the host RNA polymerase is inhibited and T7 RNA polymerase assumes responsibility for transcribing middle and late-stage genes^27,28^. T7 RNA polymerase is thus a “lynchpin” gene and we reasoned that a ligand-responsive promoter regulating this gene (*gp1*) would be an optimal site to exert systemic control. The region upstream of *gp1* gene is comprised of several parts (Fig. 1A). The beginning of the genome contains elements used for replication, including terminal repeats, promoter A0 recognized by the host RNA polymerase, and origin of replication øOL^29^. Downstream of these components lies early promoters A1, A2 and A3, all of which are recognized by the host RNA polymerase, followed by early phage genes *gp0*.*3, gp0*.*4, gp0*.*5, gp0*.*6*, and *gp0*.*7*. These genes are involved in host suppression and inhibition of host defenses^30^.

**Figure 1.**
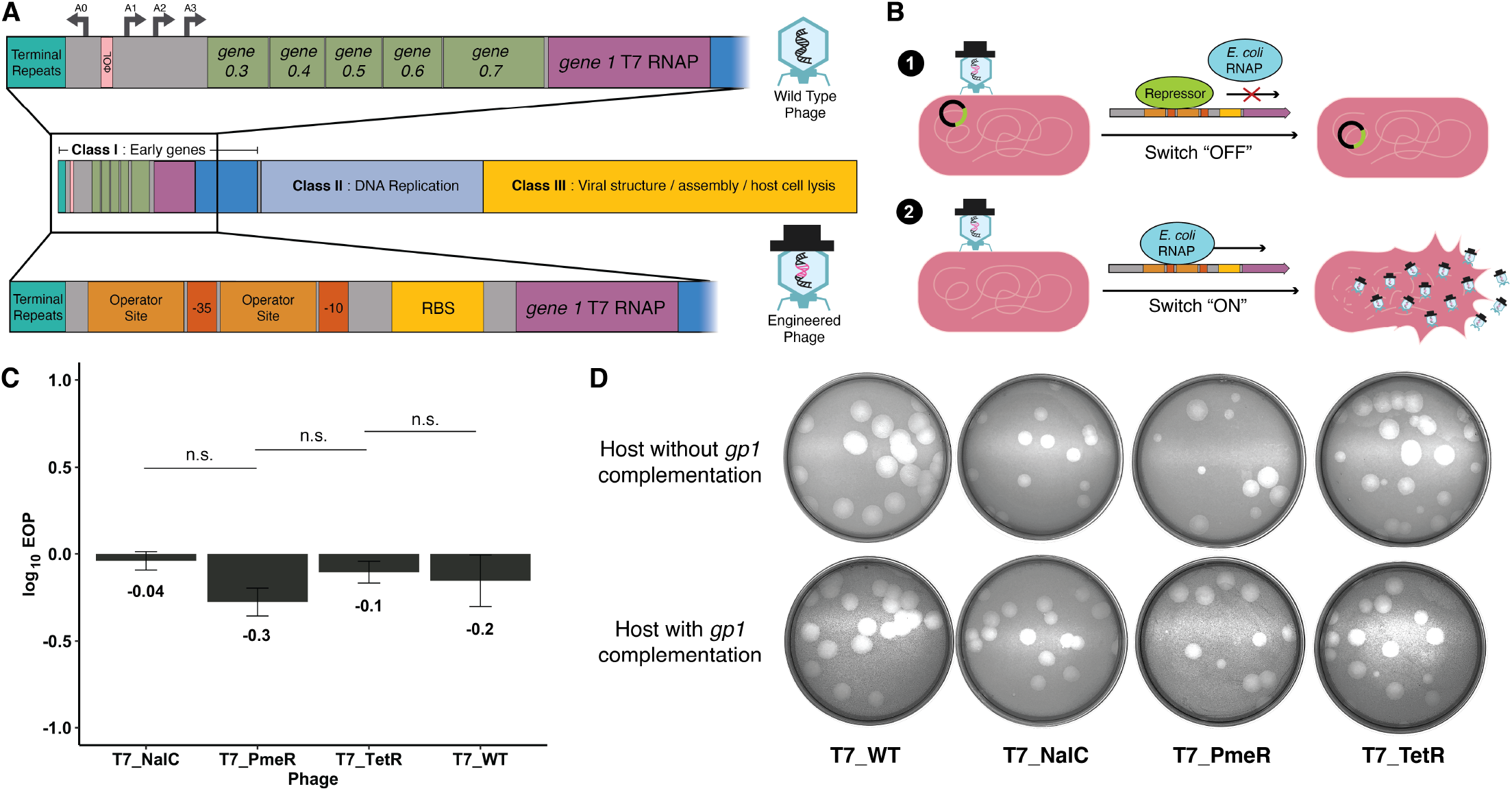
Engineered phages retain wildtype infectivity. **(A)** Schematic illustration of the T7 bacteriophage genome with early genes shown in an expanded closeup. The organization of the phage genome is shown in the middle, wildtype phage is shown top, and the engineered phage is shown bottom. The portion of the phage genome shown boxed in black is engineered with a replacement synthetic promoter and ribosome binding site (RBS). **(B)** On top (1), an illustration of the phage “OFF” state, where repressor (green) prevents RNA Polymerase (RNAP, blue) binding and blocks *gp1* transcription, stopping phages from killing host. On bottom (2), a schematic illustration of the phage “ON” state. Infection without the repressor present allows RNAP to transcribe *gp1* and phages replicate and kill the bacterial host. **(C)** Ability of engineered T7 phages to infect *E. coli* by Efficiency of Plating (EOP) using *E. coli* complementing *gp1* as a reference host. Compared to wildtype T7 (T7_WT), T7 engineered with NalC, PmeR, and TetR (T7_NalC, T7_PmeR, and T7_TetR, respectively) show no significant difference (n.s.) in ability to plaque. Data is represented as mean ± SD of biological triplicates. **(D)** Plaque morphology for wildtype and engineered phages shown by plaque assay on wildtype *E. coli* and *E. coli* with *gp1* complemented host after a 19-hour incubation. Plaques for engineered phages retain wildtype plaque morphology.

We assembled engineered T7 phage genomes in yeast without the genomic segments covering early promoters A0, A1, A2, A3, origin of replication øOL, and gene products *gp0*.*3, gp0*.*4, gp0*.*5, gp0*.*6*, and *gp0*.*7*, thereby retaining the essential replication elements but simplifying the early genome to create an effective regulatory switch. This approximately 3 kb long genomic tract was replaced with 95-100 bp long ligand-responsive promoter NalC, PmeR, or TetR (T7_NalC, T7_PmeR, and T7_TetR, respectively) placed immediately upstream of *gp1* (T7 RNA polymerase) (Fig. 1A). Engineered phages were subsequently ‘rebooted’ by transforming their genome into an *E. coli* host^31^. Once an engineered phage infects a host expressing the repressor, the repressor will bind to its cognate promoter and block the expression of *gp1*, attenuating phage activity (Fig. 1B). In the presence of the inducer or in a host where cognate repressor is absent, the repressor is unable to bind to the promoter, initiating transcription of *gp1* and reactivating the phage (Fig. 1B).

Since phage amplification occurs rapidly within the host, we wanted to evaluate different repressor-promoter systems for their OFF-states. We chose to test repressor-promoter systems NalC, PmeR, and TetR as these systems have been previously engineered with tight and inducible gene regulation^32–34^. However, before comparing the ability of different repressor-promoters to regulate the phage, we first sought to determine if our engineered phages (T7_NalC, T7_PmeR and T7_TetR) were viable after removing approximately 3 kb (∼7%) of native genes and regulatory regions. The engineered phages were viable despite the substantial genomic disruption. Efficiency of plating (EOP) assays showed no significant difference (*p-*value > 0.05) in ability to plaque between our engineered phages and T7_WT (Fig. 1C). Furthermore, the plaque morphology of the engineered phages was indistinguishable from T7_WT in *E. coli* with and without *gp1* complementation (Fig. 1D). Taken together, our engineered phages were able to maintain viability and infectivity on susceptible bacterial host compared to T7_WT.

### Repressing *gp1* diminishes activity of synthetic phages

We applied T7_NalC, T7_PmeR, and T7_TetR on *E. coli* host expressing the cognate repressor from a plasmid (Fig. 2A, *E. coli_*NalC, *E. coli_*PmeR and *E. coli_*TetR, respectively) or control wildtype *E. coli* host without repressor (Fig. 2A, *E. coli*_WT). For each engineered phage, we evaluated the estimated latent period (the time required to complete one infection cycle) by comparing cell densities at two different multiplicities of infection (MOI or the phage-to-bacteria ratio). The latent period can be estimated from the inflection in the growth curves between two MOIs where the difference is equal to the phage burst size (see Materials and Methods) (Fig. 2A). All three synthetic phages showed a delayed latent period in host expressing the cognate repressor compared to *E*.*coli*_WT lacking the repressor (Fig. 2A and 2B). The TetR repressor had the strongest attenuation of phage activity among the three repressors based on estimated lysis time (Fig. 2A). In *E*.*coli*_WT, T7_PmeR and T7_TetR required approximately 20 minutes to lyse the host, which was comparable to that of T7_WT (Fig. S1). This is also consistent with similar EOPs for wildtype and engineered phages (Fig. 1B). T7_NalC required a longer time of 30 minutes to lyse the host, indicating that the NalC promoter may be affecting phage fitness even in the absence of the repressor. When engineered phages were applied on hosts expressing their corresponding repressors, the estimated lysis time of hosts *E. coli_*PmeR, *E. coli_*NalC and *E*.*coli_*TetR was approximately 40, 30 and 40 minutes, respectively. As expected, the repressor was able to delay the activity of engineered phages (Fig. 2B). We found that the TetR system gave the longest delay (2-fold) in estimated lysis time after normalizing for differences in basal activity of each engineered phage on *E*.*coli*_WT (Fig. 2C). We also examined residual bacterial density (OD_600_) as it is indicative of the fraction of host cells that remain unlysed. After 6 hours of incubation with the engineered phages, bacterial density was highest for *E. coli_*TetR, followed by *E*.*coli_*NalC and then *E*.*coli_*PmeR (Fig. 2A). Therefore, synthetic switches engineered into the phages able to reduce phage efficacy and delay host cell lysis. Since the TetR system gave the longest delay, we chose *E*.*coli_*TetR and T7_TetR for further optimization.

**Figure 2.**
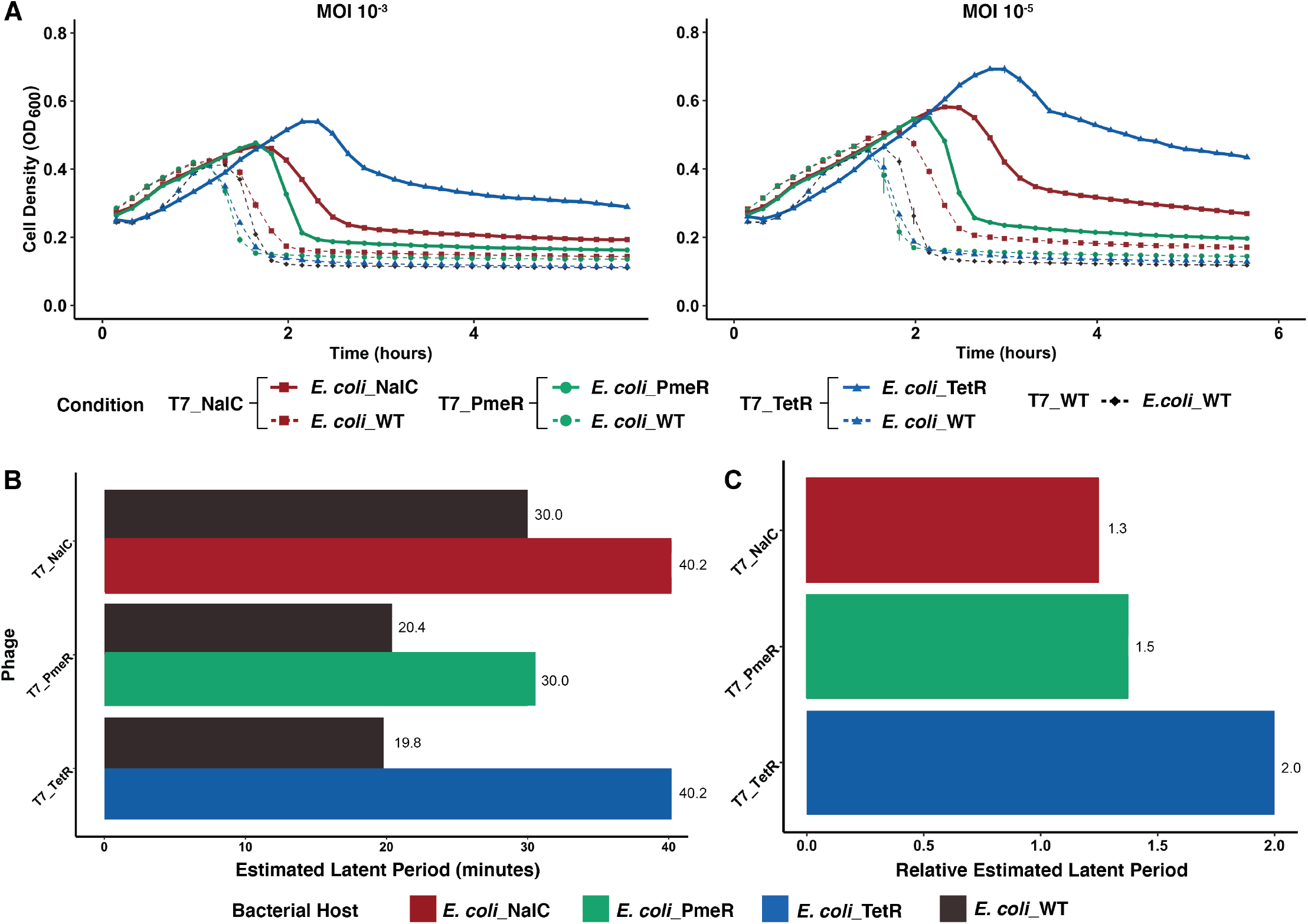
Repression of *gp1* reduces activity in engineered phages. **(A)** Bacterial cell density over time measured by absorbance (OD_600_) of wildtype host (*E. coli_*WT; dotted lines) and host expressing NalC, PmeR or TetR repressor (*E. coli_*NalC, *E. coli_*PmeR, *E. coli_*TetR, respectively; solid lines) after addition of T7_NalC (red), T7_PmeR (green) or T7_TetR (blue) repressors or T7_WT (black). Phages were applied at time 0 at a MOI of 10^−3^ and 10^−5^. Phages with synthetic promoters have delayed lysis on hosts expressing the cognate repressor but have a comparable estimated latent period in hosts without repressor. All data represented as mean OD_600_ ± SD in technical triplicates. **(B)** The estimated latent period of engineered phages on wildtype host compared to host expressing the cognate repressor, as determined by comparing the inflection in the growth curves between two MOIs where the difference is equal to the phage burst size **(C)** The relative estimated latent period of engineered phage on hosts expressing the cognate repressor compared to wildtype host. See also Figure S1.

### Engineering ribosome binding sites further enhances phage control

Although T7_TetR performed better than T7_PmeR and T7_NalC, T7_TetR still showed relatively high activity in the OFF-state (Fig. 2C). We hypothesized this could be due to high basal expression of *gp1* from the TetR-regulated promoter. To address this, we investigated if modifications to the ribosome binding site (RBS) could further attenuate phage activity in the OFF-state by reducing basal expression of *gp1*. We engineered T7_TetR with three RBS variants of the TetR-regulated promoter. The chosen RBS variants spanned a range of translational activities based on a large-scale experimental study of *E. coli* RBS variants^32,35^. Relative to the strength of the original RBS at 100% strength (T7_TetR_RBS1), we engineered T7_TetR with RBS variants at 39% strength (T7_TetR_RBS2), 3% strength (T7_TetR_RBS3), and 1% strength (T7_TetR_RBS4) (see Methods). On *E*.*coli*_WT without repressor, all T7_TetR RBS variants required an estimated 15-20 minutes to lyse the host, comparable to T7_WT, indicating that strength of RBS had no measurable impact on phage activity in the absence of repression (Fig. 3A and 3B). In contrast, when the engineered phages were applied on *E. coli_*TetR, there was substantial delay in time required to lyse the host compared to *E. coli_*WT (Fig. 3B). The time required to lyse *E. coli_*TetR was inversely related to the strength of the RBS. T7_TetR_RBS4 (the weakest RBS) required 95 minutes to lyse *E. coli_*TetR, the longest time and a 4.75-fold greater delay relative to *E. coli_*WT (Fig. 3C). T7_TetR_RBS4 also retained a high residual bacterial density after 6 hours, suggesting lowering basal expression of *gp1* successfully attenuated T7 activity (Fig. 3A). Weaker RBSs are thus able to provide enhance control of phage replication by further restricting phage activity in the OFF-state.

**Figure 3.**
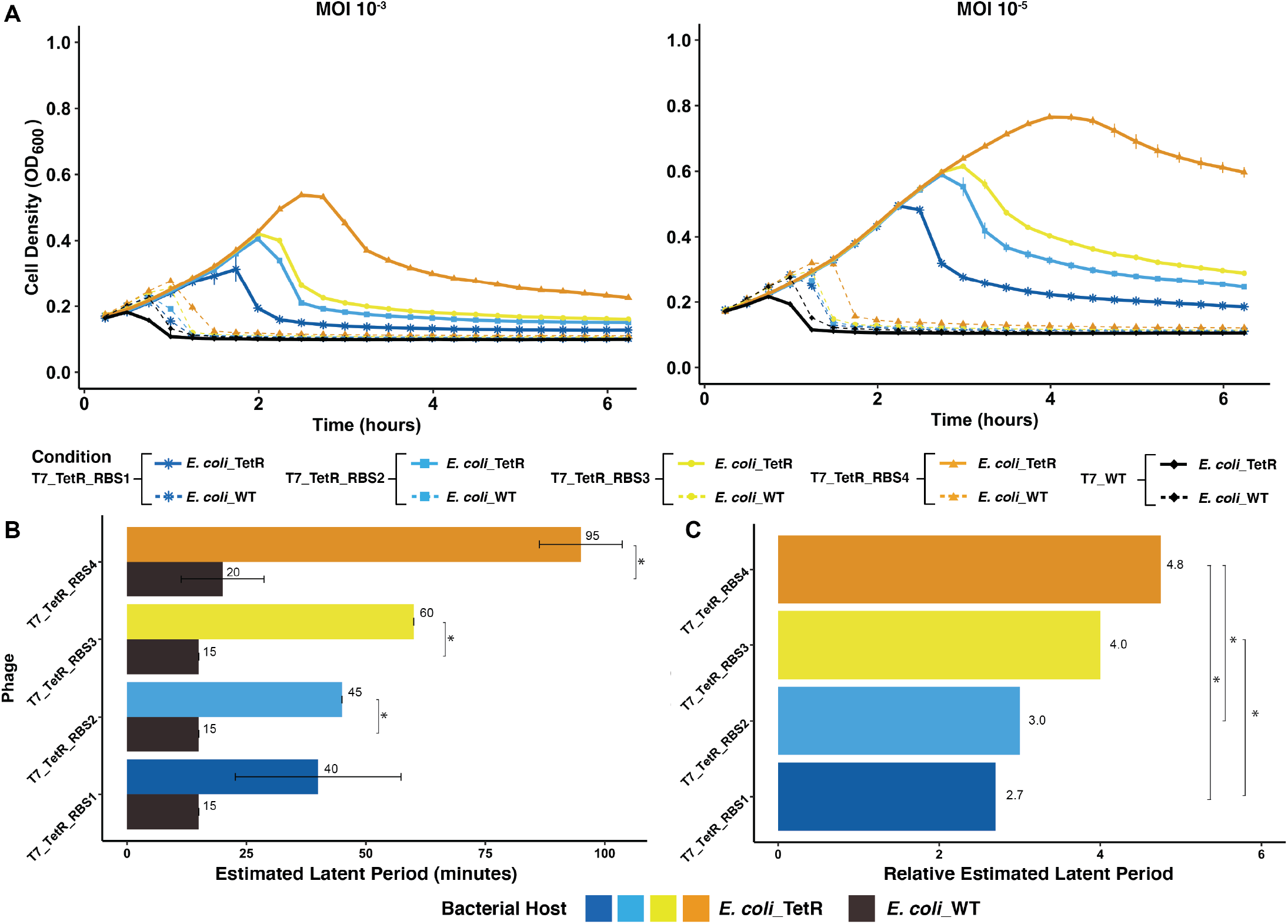
Engineered ribosome binding sites further enhances control over phage activity. **(A)** Bacterial cell density over time measured by absorbance (OD_600_) for wildtype (*E. coli*_WT; dotted lines) and host expressing TetR repressor (*E. coli_*TetR; solid lines) after application of engineered T7 with an RBS at 100% strength (T7_TetR_RBS1, dark blue), 39% strength (T7_TetR_RBS2, light blue), 3% strength (T7_TetR_RBS3, yellow) and 1% strength (T7_TetR_RBS4, orange). Phages were added at time 0 at a MOI of 10^−3^ and 10^−5^. The delay in estimated latent period increases as the RBS strength decreases. All data represented as mean OD_600_ ± SD in biological triplicates. **(B)** The estimated latent period of engineered phages on *E*. coli_WT compared to hosts expressing the TetR repressor. **(C)** The relative estimated latent period of engineered phage on host expressing the TetR receptor compared to wildtype host. Single asterisk (*) represents significant difference (*p-*value < 0.05). Non-significant difference not shown. See also Figure S2.

### Repression significantly delays replication in engineered phages

To further characterize the effectiveness of our synthetic switch, we examined the number of progeny phages over one infection cycle using one-step growth curves for T7_WT and T7_TetR_RBS4 on *E. coli_*WT and *E*.*coli_*TetR (Fig 4A). By counting the number of phages produced at different times, we determined the actual latent period (the time required to complete one infection cycle), and compared the log change in phage titer over one infection cycle. In prior literature, the latent period for T7_WT is approximately 20 minutes with an expected 2-2.5 log increase in the total phage population after one infection cycle^36,37^. Since the TetR promoter and RBS does not have an impact on T7 fitness, we expected similar values for T7_TetR_RBS4 on *E. coli_*WT.

**Figure 4.**
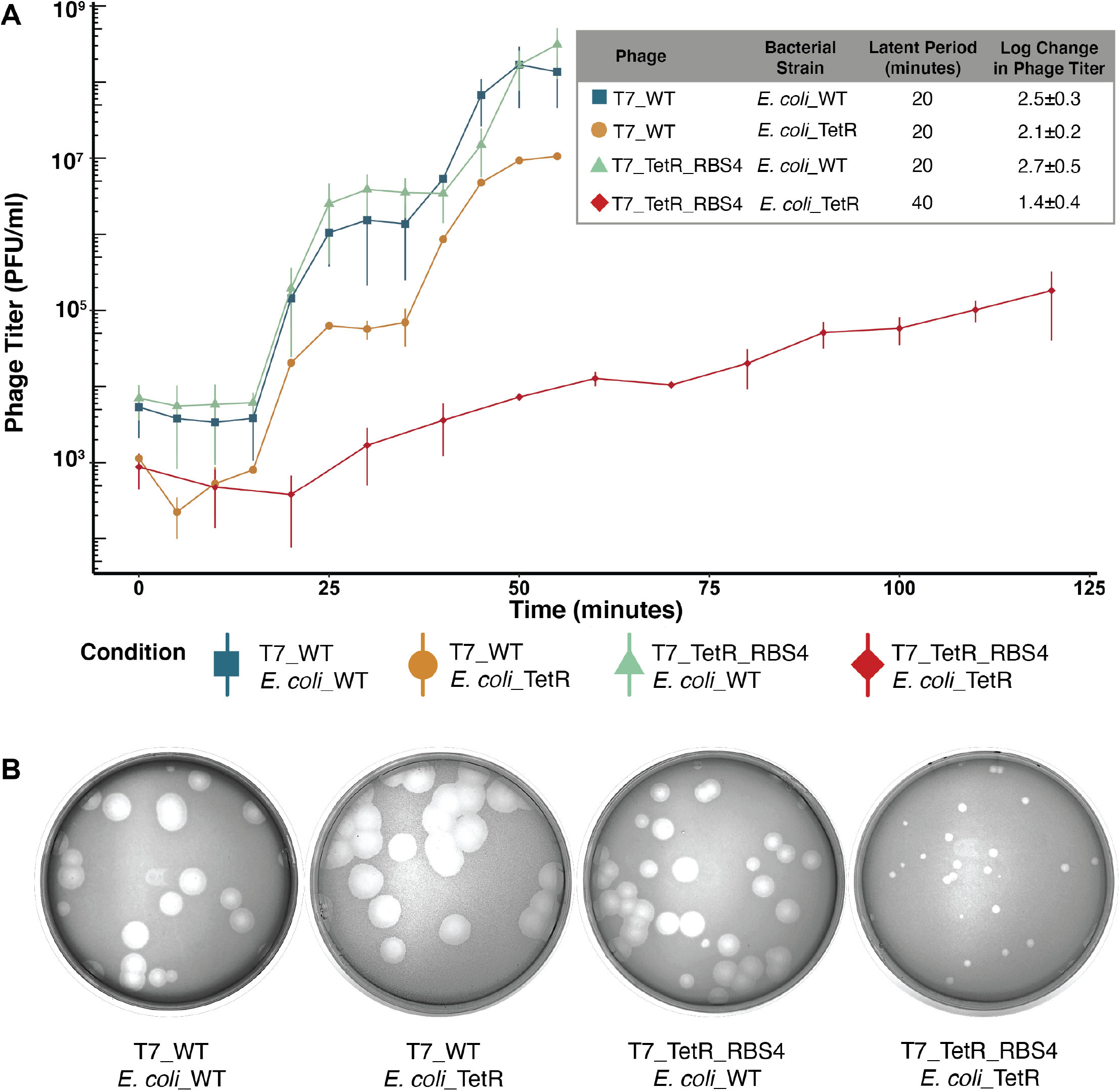
Repression significantly delays replication in engineered phages. (A) One step growth assay showing phage titer (PFU/ml) of wildtype (T7_WT) and engineered (T7_TetR_RBS4) phage on wildtype (*E. coli_*WT) and engineered (*E. coli_*TetR) host. Inset table shows the latent period (minutes) and an estimated log change in phage titer produced from the first replication cycle. All data shown as mean ± SD in biological triplicates. **(B)** Plaque assay of wildtype (T7_WT) and T7 with TetR and a 1% strength ribosome binding site (T7_TetR_RBS4) phages on wildtype host (*E. coli_*WT) and host expressing TetR repressor (*E. coli_*TetR) host after 19 hours of incubation.

Our one step assay confirmed that both T7_WT and T7_TetR_RBS4 have a 20 minute latent period on *E*.*coli_*WT and T7_WT has a 20 minute latent period on *E*.*coli_*TetR (Figure 4A). After one infection cycle, the average increase in total phages for T7_WT on *E. coli_*WT and *E. coli_*TetR was approximately 2.5±0.3 and 2.1±0.2 log respectively, while T7_TetR_RBS4 saw a comparable increase in total phages of 2.7±0.5 log on *E. coli*_WT. These results confirm that the TetR repression system had no apparent effect on phage activity for T7_WT or for T7_TetR_RBS4 in the absence of repression. In contrast, T7_TetR_RBS4 infecting *E. coli*_TetR had greatly extended latent period of 40 minutes with only a 1.4±0.4 log increase in total phages (Fig. 4A), indicated a dramatic delay of phage activity with very gradual production of phage progeny during each replication cycle. We next determined how the TetR_RBS4 repression system affects the ability of the phage to plaque on *E. coli*_TetR compared to *E. coli_*WT using an EOP assay. T7_TetR_RBS4 phages had an EOP of -0.2±0.1 while T7_WT phage had an EOP of 0.2±0.1. The plaque activity of T7_TetR_RBS4 and T7_WT was slightly significantly different (*p-*value = 0.05) and the T7_TetR_RBS4 plaques were significantly smaller than T7_WT plaques after nineteen hours of incubation, indicative of much slower phage activity and consistent with our one-step results (Fig 4B). In summary, engineered T7_TetR_RBS4 phages have a significant delay in replication compared to wildtype when the synthetic repressor is present, and phage activity can be fully recovered if the synthetic repressor is not present or under induction.

### Phage infectivity can be further controlled by tuning inducer concentration

To assess if phage activity could be dynamically controlled using a small molecule inducer, we compared the activities of T7_TetR_RBS4 with and without the inducer anhydrotetracycline (aTC)^38^. Comparison of spot plates showed that repressed T7_TetR_RBS4 (Fig. 5A, left) regained activity comparable to T7_WT when maximally induced at 1 μM aTC (Fig 4A, right, Fig. S2). A control experiment confirmed that aTC had no impact on activities of T7_TetR_RBS4 on *E. coli_*WT host (Fig. S3). However, host cells grown over a gradient of aTC concentrations experienced a minor fitness deficit at higher aTC concentrations suggesting inducer toxicity may have partially contributed to cell death^39^. To evaluate if the activity of T7_TetR_RBS4 could be tuned by the inducer in a dose-dependent manner without cell toxicity, we performed a time course experiment over a range of aTC concentrations from 0 to 1 μM (Fig. 5B). T7_TetR_RBS4 showed switch-like behavior from no activity to high activity with modest dose-dependent activity over a narrow range of aTC concentration from 15-23 nM. Above 23 nM, T7_TetR_RBS4 was fully switched ‘ON’ and activity was comparable to activity at maximal induction at 1 μM (Fig. 5B). Below 15 nM, the activity of T7_TetR_RBS4 was nearly the same as the activity with no inducer. The tunable concentration range (15-23 nM) showed large variations which arise due to high stochasticity in the expression of *gp1* across the host population at these concentrations. Since T7 RNA polymerase can be recycled for phage gene expression^40^, even stochastic bursts of expression would eventually lead to bacterial lysis. Altogether these results show that a moderate aTC concentration can provide a tunable level of control over phage activity.

**Figure 5.**
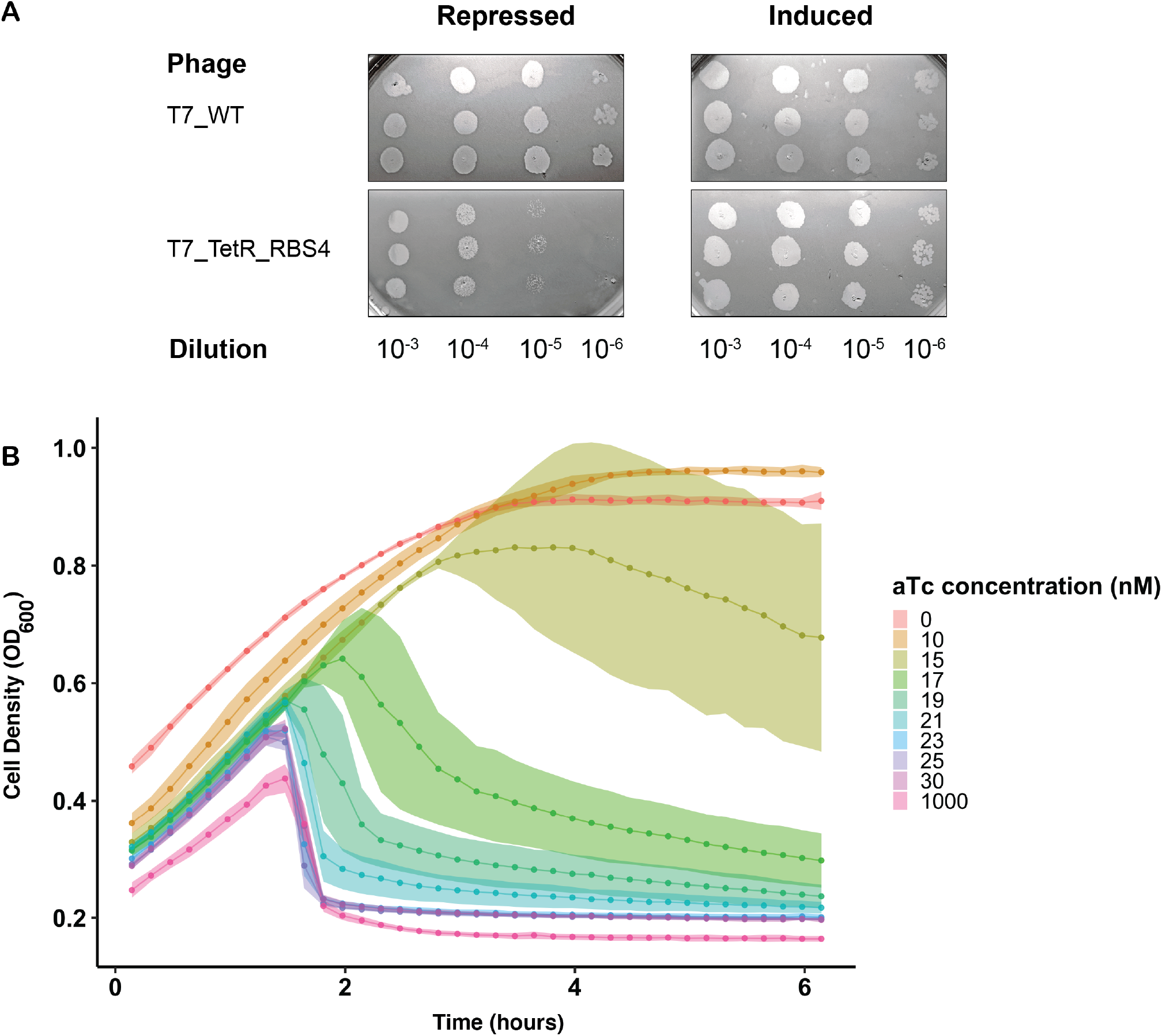
Phage infectivity can be tuned by changing inducer concentration. **(A)** Spot plate assay measuring the ability of wildtype T7 (T7_WT) and T7 with TetR and a 1% strength ribosome binding site (T7_TetR_RBS4) to infect *E. coli* expressing the TetR repressor without inducer (Repressed) or with 1 μM anhydrotetracycline (aTC) inducer (Induced). Phages are spotted from 10^−3^ to 10^−6^ dilution and spots are imaged after 4 hours of incubation. Induction restores the phage ability to spot comparable to wildtype. **(B)** Bacterial cell density over time measured by absorbance (OD_600_) for *E. coli* expressing TetR repressor. T7_TetR_RBS4 phage is added at timepoint 0 at a MOI of 10^−5^, and each culture contains aTC inducer ranging from a concentration of 0 to 1000 nM (color gradient). See also Figure S3.

## Discussion

In this study, we engineered synthetic gene regulation into T7 phage to create a phage-bacterial system with ligand-regulated infectivity. We deleted a large tract of the genome from T7 phage and inserted a suite of ligand-regulated promoters to control the expression of *gp1*, the T7 RNA polymerase. The deletion of phage early promoters and genes and replacement with ligand-regulated promoter resulted in a viable phage with infectivity comparable to T7_WT. Our most optimized engineered phage, T7_TetR_RBS4, had a 2-fold delay in estimated latent period and an approximately 2-fold decrease in change in phage titer after one replication cycle compared to T7_WT. The activity of the engineered phage could be fully recovered in dose-dependent manner by adding the inducer ligand.

Our engineered phage sets the stage for several improvements to further refine and control phage infectivity. Because the infectivity of our engineered phage was determined solely by control over T7 RNAP, even basal levels of expression would allow the phage to escape repression proceed with infection. Adding multiple levels of repression throughout the phage genome could substantially improve our system and decrease the activity the phage in the OFF state^41,42^. Critical genes such as late structural genes would make natural targets for improving control over the phage replication. Our study and other previous reports suggest that the phage genome may be tolerant to such changes without affecting viability^36,43^.

An engineered phage with a dynamic controllable infectivity switch can be used as a potent tool for bacterial community control^44,45^. Utilizing different repression systems across different bacterial hosts, phages can be redirected to different bacteria in the same community^46^. Phages could also be continuously propagated in a community using a ‘feeder’ host whose susceptibility could be turned ON or OFF as needed. By tuning inducer concentrations^47,48^, phages could be engineered to repress but not eliminate specific bacterial hosts in a complex community. Alternatively, our engineered phages can act as pseudo-lysogenic phages, maintaining a low burden on the targeted microbe until being triggered to eliminate the host. This could be useful for applications like starter cultures, where timed bacterial lysis is a critical consideration for cheese maturation^49^. Our study presents a simple genetic regulatory technique that can be further engineered to create a more controllable phage-bacterial system to precisely manipulate microbial communities^50–52^.

## Supporting Information

**Figure S1. Expression of repressors has no effect on wildtype T7**

Bacterial cell density over time measured by absorbance (OD_600_) of wildtype T7 (T7_WT) on wildtype host (*E. coli_*WT; black) and host expressing NalC (*E. coli_*NalC, red), PmeR (*E. coli_*PmeR, green) or TetR (*E. coli_*TetR, blue) repressor. Phages were applied at time 0 at a MOI of 10^−3^ and 10^−5^. There is no delay of estimated latent period when any repressor is expressed compared to wildtype host. All data represented as mean OD_600_ ± SD in technical triplicates.

**Figure S2. Activity of engineered phages is rescued by aTC inducer**

Bacterial cell density over time measured by absorbance (OD_600_) for *E. coli* expressing TetR repressor after application of no phages (T7_None, grey), wildtype phage (T7_WT, black), engineered T7 with an RBS at 100% strength (T7_TetR_RBS1, dark blue), 39% strength (T7_TetR_RBS2, light blue), 3% strength (T7_TetR_RBS3, yellow) and 1% strength (T7_TetR_RBS4, orange). Phages and 1 μM aTC was added at timepoint. Addition of aTC rescues phage function and results in comparable time to lysis for engineered phages and T7_WT. All data represented as mean OD_600_ ± SD in biological triplicates.

**Figure S3. Inducer has no effect on bacterial growth or phage activity**

Bacterial cell density over time measured by absorbance (OD_600_) for **(A)** *E. coli* expressing TetR (*E. coli_*TetR) without phage (T7_None), **(B)** wildtype *E. coli* (*E. coli_*WT) and engineered phage with a 1% strength RBS (T7_TetR_RBS4), and **(C)** *E. coli* expressing TetR (*E. coli_*TetR) and wildtype phage (T7_WT). Anhydrotetracycline (aTC) is added at concentrations from 0-1000 nM (color gradient) at time zero. *E. coli* is able to grow productively under all aTC concentrations and application of phages results in a reduction of cell density at the same time for all aTC concentrations. All data represented as mean OD_600_ ± SD in biological triplicates.

**Table S1: Primers and Plasmids used in this study**

**Table S2: Promoter and Ribosome Binding Site sequences**

## Materials and Methods

### Microbes and Culture Conditions

*Escherichia coli* (*E. coli*) 10G, a highly competent derivative of DH10β was obtained from Lucigen (60107-1)^53^, T7 bacteriophage was obtained from ATCC (ATCC^®^ BAA-1025-B2), and *Saccharomyces cerevisiae* BY4741^54^ is a laboratory stock.

Bacterial cultures were grown in LB (Luria-Bertani) media (1% Tryptone, 0.5% Yeast Extract, 1% NaCl). For plating, LB agar contains 1.5% agar, while top agar contains 0.5% agar (Teknova). Spectinomycin (115 μg/ml final concentration, GoldBio^®^) was added to media for selections of pSC101_NalC, pSC101_PmeR and pSC101_TetR. All incubations of bacterial and bacteriophage cultures were performed at 37°C, with the liquid culture shaking at 200-250 rpm consistently, otherwise specified.

T7 bacteriophage was grown and propagated using *E. coli* 10G in LB media. Phage stocks were tittered using plaque assay and stored in LB at 4°C.

*S. cerevisiae* BY4741 was grown in YPD (2% Peptone, 1% Yeast Extract, 2% Glucose) media prior to transformation. Yeast transformants were selectively grown on SD-Leu (0.17% Yeast Nitrogen Base, 0.5% Ammonium Sulfate, 0.162% Amino Acid-Leucine, [Sigma Y1376], 2% Glucose). YPD and SD-Leu plates contain 2.4% and 2% agar additionally, respectively. Yeast incubation was performed at 30°C, with liquid culture shaking at 200-250 rpm.

Short-term storage of liquid culture and plates were performed at 4°C and long-term storages of bacterial and yeast stock culture were performed at -80°C in screw-capped cryotubes, with 25% glycerol added as a cryoprotectant.

Bacterial and phage transformants were recovered in SOC (2% Tryptone, 0.5% Yeast Extract, 0.2% 5 M NaCl, 0.25% 1 M KCl, 1% 1 M MgCl_2_, 1% 1 M MgSO_4_, 2% Glucose) liquid media.

### General Cloning Procedure

PCR and cloning were adapted and performed using standard laboratory procedures^12^. Briefly, PCR amplification was performed using KAPA HiFi (Roche KK2101) for all amplifications with plasmid or phage templates. KAPA2G Robust PCR kits (Roche KK5005) were used to perform colony PCR and multiplex PCR for a screening of Yeast Artificial Chromosomes (YACs). All primer oligos were obtained from IDT™. Golden Gate assembly was performed using New England Biosciences (NEB) Golden Gate Assembly Kit (BsaI-HFv2, E1601L). DNA purification was performed using EZNA Cycle Pure Kits (Omega Bio-tek D6492-01) using centrifugation protocol. YAC extraction was performed using YeaStar Genomic DNA Extraction Kits (Zymo Research D2002). Gibson Assembly mixture was made in the laboratory (final concentration 100 mM Tris-HCL pH 7.5, 20 mM MgCl_2_, 0.2 mM dATP, 0.2 mM dCTP, 0.2 mM dGTP, 10 mM dTT, 5% PEG-8000, 1 mM NAD^+^, 4 U/ml T5 exonuclease, 4 U/μl Taq DNA Ligase, 25 U/mL Phusion polymerase). PCR product visualization was performed using agarose gel electrophoresis with appropriate agarose concentration and SYBR^®^ Safe DNA Gel Stain (Invitrogen).

PCR amplification using plasmid templates was performed with 0.1 ng DNA template. Phage fragment amplification was performed using 1 μl phage crude lysis treated at 65ºC for 10 minutes as a template. Deletions and insertions of the T7 genome were performed using PCR primers skipping or adding desired sequences, respectively. All plasmid-template PCR products were treated with DpnI (NEB) following standard protocol. Briefly, purified PCR product was combined with 5 μl 10x CutSmart Buffer, 1 μl DpnI, and dH_2_O to 50 μl reaction. Digestion was performed at 37°C for at least 2 hours, followed by heat inactivation at 80°C, 20 minutes. PCR purification was performed afterward. All PCR products were quantified using NanoDrop 2000 Spectrophotometer (Thermo Scientific).

*E. coli* 10G competent cells were made by mixing 192 mL SOC with 8-ml overnight culture and incubating at 21°C and shaking at 200 rpm until OD_600_∼0.4 was reached, determined by the Agilent Cary 60 UV-Vis Spectrometer using manufacturer documentation. Cells were centrifuged at 4°C, 800 xg to1000 xg for 20 minutes, the supernatant was discarded, and cells were resuspended in 50 ml of pre-cooled 10% glycerol. Centrifugation and washing were repeated three times. Cells were resuspended in a final volume of ∼1 ml 10% glycerol and were aliquoted and stored at - 80°C. Cells made by this protocol are competent for both plasmid and YAC transformation using electroporation.

Electroporation was performed using 40 μl 10G competent cell for both plasmids and YACs using a Biorad MicroPulser (165-2100), EC2 setting with 2-mm cuvette, 2.5 kV, single pulse. All cuvettes and Eppendorf tubes were chilled prior to the electroporation. After electroporation, recovery was performed by adding 950 μl pre-warmed SOC and incubated at 37°C for 1 hour and plated on relevant selective media.

Detailed protocols for cloning are available on request. All primers and plasmids used in this study are listed in supporting document (Table S1).

### Phage Engineering

A large segment (2961 bp) in the left end region of the wild type (WT) T7 phage genome, including promoters A0, A1, A2, A3 and *gp0*.*3-0*.*7* genes, was deleted. Two identical operator (repressor binding) sites were inserted, one upstream of -35 and the other in between the -35 and -10 consensus sequences replacing the spacer sequence of promoter A1 driving the expression of *gp1* gene. The three operator sites are NalC1, PmeR2, and TetO correspond to repressors, NalC, PmeR and TetR, respectively, in bacterial system. The Bujard RBS (Ec-TTL-R111) in TetO engineered phage was replaced by BBa_J61106 (Ec-TTL-R065), BBa_J61133 (Ec-TTL-R003), or DeadRBS (Ec-TTL-R001) with 39%, 3%, and 1% translational strength relative to the Bujard RBS as 100% strength^35^, respectively. The sequences of all consensus sequences, operator sites, RBS, and repressors used are included in supporting document (Table S2).

Engineered phage genomes were assembled using yeast assembly^4,5^, which requires yeast transformation of relevant DNA segments. A prs315 yeast centromere plasmid was split into three segments by PCR, separating the centromere and leucine selection marker, which has been shown to improve assembly efficiency by limiting recircularization events^6^. T7 genomic segments were made by PCR using WT T7 as a template.

Relevant DNA fragments were mixed (0.1 pmol/fragment) and transformed into *S. cerevisiae* BY4741 using the high efficiency yeast transformation protocol^7^. Successfully assembled YACs were selected using SD-Leu media. After 2-3 days of incubation at 30°C, colonies were picked and directly assayed by multiplex colony PCR to screen for colonies with correctly assembled YACs. Multiplex PCR was an effective way of distinguishing correctly assembled YACs by interrogating junctions in the YACs. Correctly assembled YACs were purified and transformed into *E. coli* 10G cells, and after 1-hour recovery, 400 μl was inoculated in 4.6 ml LB. This culture was incubated until a complete lysis, engineered phages were purified, sequence-confirmed, and stored at 4°C in LB.

### Plasmid Descriptions

pSC101_NalC, pSC101_PmeR and pSC101_TetR contain a pSC101 backbone, spectinomycin resistance cassette, and repressors NalC, PmeR and TetR under constitutive promoter apFAB61, respectively.

pHT7_gp1 contains a pBR backbone, kanamycin resistance cassette, mCherry fluorescence marker, and the T7 RNAP gene, *gp1*. Both mCherry marker and *gp1* are under constitutive expression.

### Bacterial Methods and Phage Titer Quantification

Bacterial concentration was determined using a drop plate with 10-fold serial dilution of bacterial culture. Ten-microliters of bacterial dilution were dropped on LB plate in triplicate and counted after overnight incubation as colony-forming unit per milliliter (CFU/ml). Optical density (OD) measurement for all microplate-reader experiments were performed using microplate reader (Synergy HTX) at 600 nm wavelength, except the preparation of electrocompetent cell and the determination of log phase of bacterial culture that were performed using an Agilent Cary 60 UV-Vis Spectrometer.

Bacterial culture used for bacteria-phage interaction experiment and phage quantification was obtained at log phase when the initial phage was added, unless otherwise specified. Briefly, overnight bacterial culture was diluted 1:20 in liquid LB media. Bacterial culture was collected when the OD_600_ reaches 0.4-0.6 as determined using an Agilent Cary 60 UV-Vis Spectrometer.

Phage stock was produced by a complete infection of phage in *E. coli* 10G. Bacterial lysate was centrifuged at 4,400 xg for 5 minutes and supernatant was filtered through a 0.22 μm filter (Celltreat 151205-051). Phage titer was determined by plaque assay with a 10-fold serial dilution of phage lysate. Typically, 10-20 μl of phage dilution is mixed thoroughly with 250 μl log-phase bacterial culture and 3.5-ml 0.5% top agar and plated on 37°C pre-warmed LB plate. Plaque number was counted after overnight incubation as plaque-forming unit per milliliter (PFU/ml). Spot assay was considered as a preliminary quantification of phage titer, which was performed occasionally prior to whole-plate plaque assay. Log-phase bacterial culture was mixed with 3.5-ml top agar and plated on pre-warmed LB plate. One-and-a-half microliters of phage dilution from 10^0^ to 10^−8^ were spotted in triplicate on set top agar. The number of PFU was approximated determined at 4-6 hours of incubation and the appropriate dilution was chosen to perform whole-plate plaque assay.

Multiplicity of Infection (MOI) was determined by dividing phage titer by bacterial concentration. MOI used in this study was approximately 10^−3^ and 10^−5^, except for one-step growth curve which was performed with MOI ∼ 0.1.

Efficiency of Plating (EOP) was calculated using wildtype *E. coli* 10G as a reference host. After performing a plaque assay using desired bacterial hosts and phages, EOP was determined by dividing the experimental phage titer to the phage titer on the reference bacterial host. Typically, a 10-fold serial dilution was performed with phage stock in triplicate and 10 μl appropriate phage dilutions were applied for plaque assay as described in desired bacterial host.

### Infection time course curves

Preliminary change in phage activity of engineered phages was determined by evaluating bacterial growth after addition of different phages. Seven microliters of overnight bacterial culture were inoculated in 133 μl of relevant media as a 1:20 dilution with corresponding antibiotics if necessary or 1 μM aTC to the total volume of 140 μl in a 96-well plate. Ten microliters of phages were added in triplicate at an MOI of 10^−3^ and 10^−5^. The culture was continuously incubated in microplate reader (Synergy HTX) at 37°C with OD_600_ measured every 10 minutes for 6 hours. The bacterial growth curve was constructed by averaging OD_600_.

### Phage latent period estimation with infection time courses

With the assumption that T7 phage produces an average of 100 progeny phage at 37°C when infecting 10G^37^, time to lysis was estimated by comparing the inflection point of bacterial growth (or the point at which the culture begins to lyse) for two cultures after addition of two phage titers spanning 2 orders of magnitude. An MOI of 10^−3^ and 10^−5^ were chosen for this estimation. The estimated latent period was calculated as the difference between the last time point immediately before the inflection point in OD_600_ between the two cultures.

### Phage Growth Quantification: Latent Period and Phage Titer change estimation

One-step phage growth assays were performed to construct one-step growth curves using an adaptation of a standard protocol^55,56^. Briefly, Seven-hundred-fifty microliters of an overnight bacterial were added into 15-ml LB media with antibiotics if necessary. The culture was incubated shaking at 250 rpm, 37°C in a 50-ml flask. Bacterial culture was collected when the OD_600_ ∼ 0.25 was reached in 15-ml tube and concentrated into 1.5 ml by centrifugation at 4,400 xg for 5 minutes. Bacterial culture was transferred into a 2-ml Eppendorf tube, phage was added into the culture at MOI ∼ 0.1. Phage-bacteria culture was incubated for absorption without shaking at 37°C for 5 minutes. Culture was washed four times by centrifugation at 10,000 xg for 30 seconds, discarding the supernatant, and resuspending with 1-ml LB media, with antibiotic if necessary. In between washes, culture was resuspended by vortexing. After the final wash, 1-ml media was added and transferred into 14-ml pre-warmed media in a 50-ml flask. Eight-hundred microliters of culture was collected immediately and centrifuged at 10,000 xg for 30 seconds. Seven-hundred-microliter supernatant was filtered through 0.22 μm (Celltreat 151205-051) into a new Eppendorf tube. Fifteen-milliliter culture was incubated shaking at 250 rpm at 37°C. The culture sample was collected every 5 or 10 minutes, depending on the type of phage culture for 60 or 120 minutes. Phage titer was quantified by plaque assay using wildtype *E. coli* 10G. Plated culture was incubated at 37°C for 12 hours and the numbers of plaques were counted and calculated. The dilution of the phage lysate was made if necessary. The growth curve of each bacteria-phage combination was constructed in biological triplicates.

The latent period of each phage-bacteria combination was quantified by the first phage replication cycle^56^ using one-step phage growth curve. The mid-timepoint at exponential-phase of replication cycle in between plateau phage titer was considered as a latent period.

The change in total phage titer was estimated using the phage growth curve by calculating the log difference of number of phage progenies at the plateau after the first replication cycle and at the initial plateau where phage titer added into the experimental culture.

### Inducer dependency assay

The activity of phage in response to a range of aTC inducer concentrations was determined by quantifying bacterial growth using OD_600_. Briefly, 7 μl overnight bacterial culture was inoculated in 133 μl of relevant medium as 1:20 dilution to the total volume of 140 μl in a 96-well plate. Media used was prepared with a range of aTC concentration from 0 to 1 μM. Culture was incubated in a microplate reader (Synergy HTX), and OD was measured at 600 nm wavelength. When bacterial OD_600_ ∼ 0.25 was reached, 10 μl of desired phage or LB as control was added into the culture for MOI ∼ 10^−5^. Culture was continuously incubated for 7 hours using vertical shaking mode, and OD_600_ was measured every 10 minutes. The experiment was performed in biological triplicates.

### Statistical Analysis

All data analyses were performed in Microsoft Excel 2020 and R v4.0.4^57^. Bonferroni pairwise *t*-test^58,59^ was used to detect the difference amongst conditions in all experiments. Statistical analyses were considered significant at *p* < 0.05.

## Author Information

### Authors

**Chutikarn Chitboonthavisuk** – *Department of Biochemistry, University of Wisconsin – Madison, 433 Babcock Dr, Madison, Wisconsin 53706, United States*

**Chun Huai Luo** – *Department of Biochemistry, University of Wisconsin – Madison, 433 Babcock Dr, Madison, Wisconsin 53706, United States*

**Phil Huss** – *Department of Biochemistry, University of Wisconsin – Madison, 433 Babcock Dr, Madison, Wisconsin 53706, United States*

**Mikayla Fernholz** – *Department of Biochemistry, University of Wisconsin – Madison, 433 Babcock Dr, Madison, Wisconsin 53706, United States*

## Author Contributions

C.C. and C.H.L. contributed equally. S.R., C.C., C.H.L., P.H., and M.F. conceptualized and designed the study. C.H.L. and M.F. constructed plasmids. C.C. and C.H.L. performed the experiment, curated the data, performed the analysis, wrote the original manuscript, and designed the figures. S.R., C.C., C.L., P.H., and M.F. reviewed and edited the manuscript. All authors discussed the results and provided critical feedback on the manuscript.

## Notes

S.R. is on the scientific advisory board of MAP/PATH LLC. All other authors declare no competing financial interest.

## Acknowledgements

This work was supported by National Institute for Allergy and Infectious Disease grant 1R21AI156785-01 (to S.R). C.C was supported by a graduate training scholarship from the Anandamahidol Foundation (Thailand).

