## Supplementary material for "Engineering a dynamic, controllable infectivity switch in bacteriophage T7": Figure S

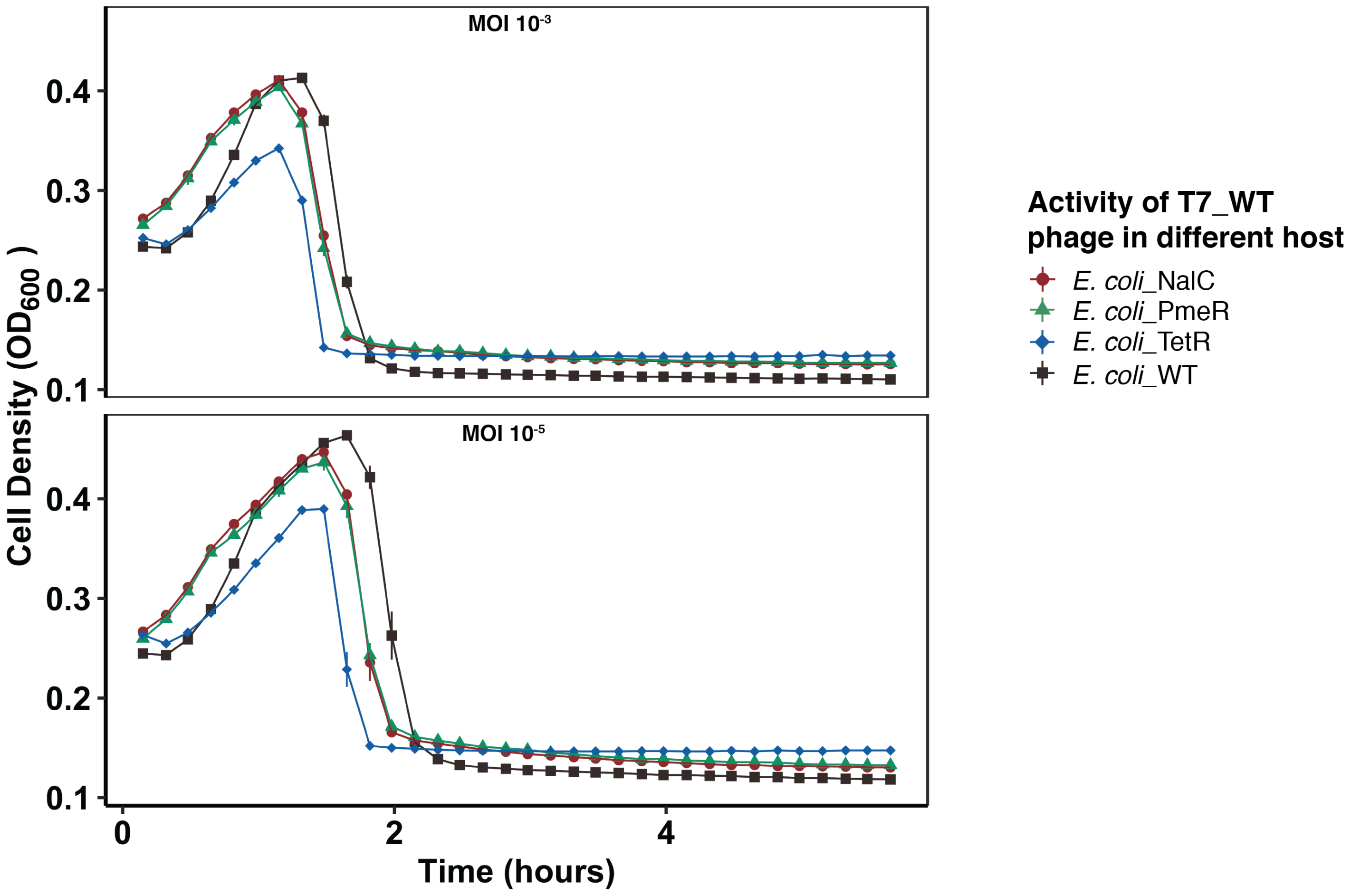


**Figure S1. Expression of repressors has no effect on wildtype T7**

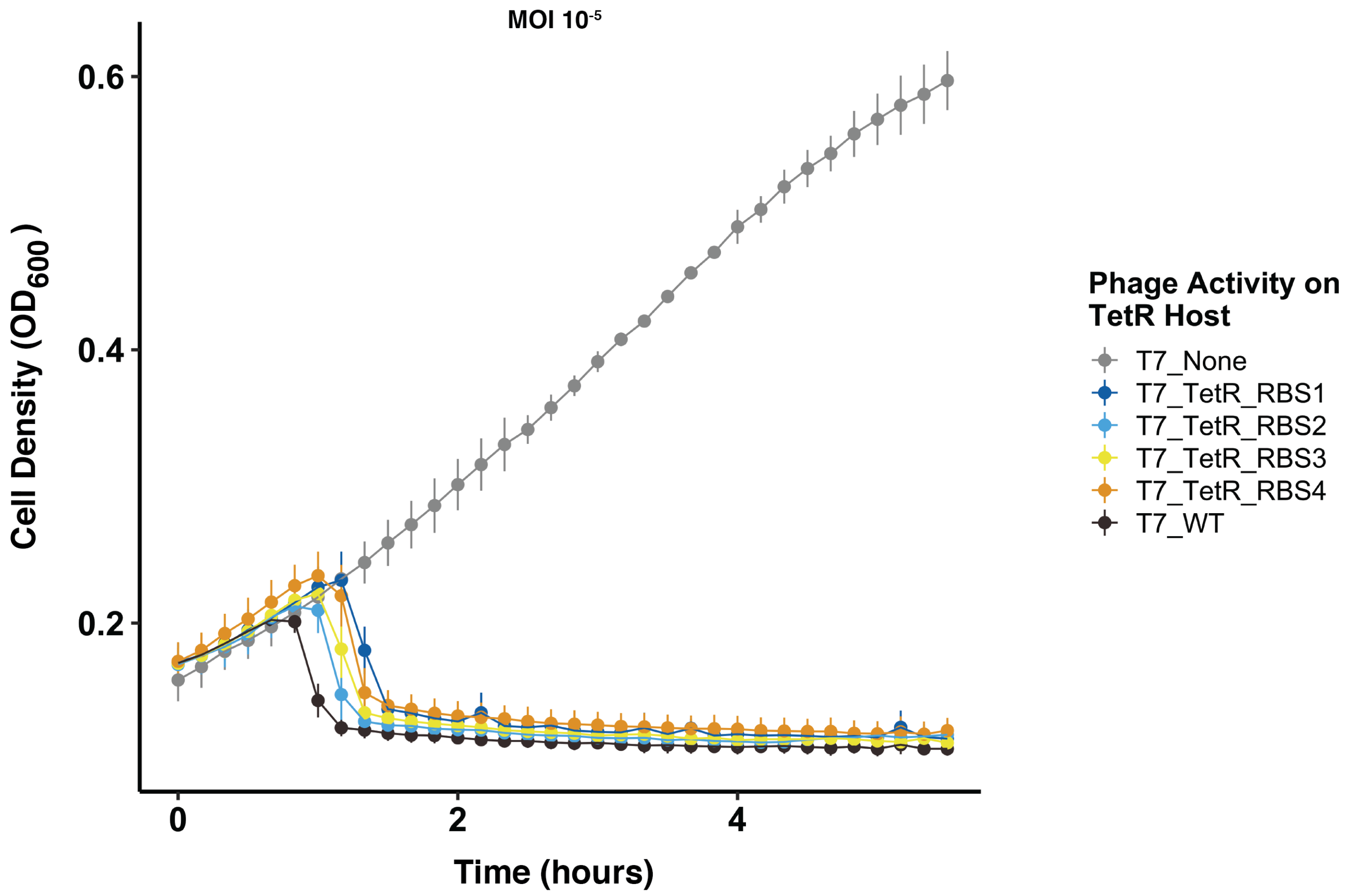


**Figure S2. Activity of engineered phages is rescued by aTC inducer**

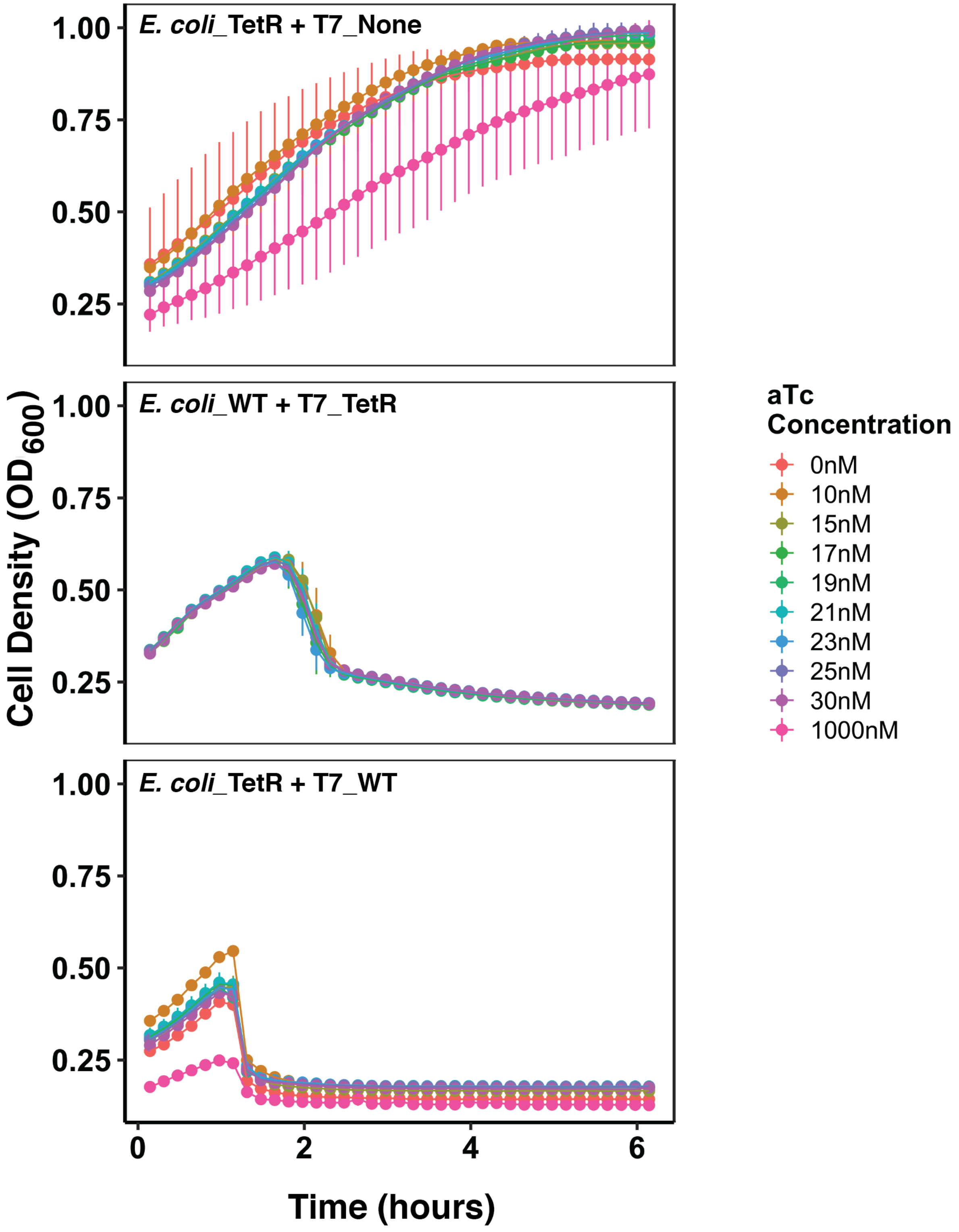


**Figure S3. Inducer has no effect on bacterial growth or phage activity**
